# Clever-1 blockade disrupts lipid metabolism and mitochondrial fitness in acute myeloid leukemia

**DOI:** 10.64898/2026.08.12.744345

**Authors:** Arno Ylitalo, Jesper Mickos, Milja Hakoniemi, Rita Turpin, Stuart Prince, Maija Hollmén

## Abstract

Therapy resistance in acute myeloid leukemia (AML) is linked to metabolic plasticity and mitochondrial fitness of leukemic stem and progenitor cells. Clever-1 is a scavenger receptor with established immunoregulatory functions, but its leukemia cell–intrinsic roles remain unclear. Here we identify Clever-1 as a regulator of mitochondrial integrity and lipid-dependent oxidative metabolism in AML. Using the anti–Clever-1 antibody bexmarilimab, we show that Clever-1 inhibition induces early mitochondrial transcriptional reprogramming, followed by suppression of oxidative phosphorylation (OXPHOS) in AML cell lines. Immunoelectron microscopy demonstrates mitochondrial localization of Clever-1, while proteomic analyses reveal altered association with mitochondrial-linked proteins, including ATAD3. Functionally, Clever-1 inhibition reduces mitochondrial delivery of lipoprotein-derived lipids, resulting in selective changes in mitochondrial lipid composition. These changes are accompanied by impaired respiratory complex IV assembly, disrupted cristae architecture, accumulation of dysfunctional mitochondria, and reduced spare respiratory capacity. AML models with high baseline OXPHOS activity are particularly sensitive to Clever-1 inhibition, with mitochondrial dysfunction exacerbated under lipid-restricted or metabolically stressful conditions. Together, these findings define Clever-1 as a regulator of mitochondrial bioenergetic resilience and a targetable metabolic vulnerability in AML.

## Introduction

Acute myeloid leukemia (AML), myelodysplastic syndromes (MDS), and chronic myelomonocytic leukemia (CMML) are clonal myeloid neoplasms characterized by disordered hematopoiesis and poor clinical outcomes driven by therapy resistance and disease recurrence or progression ^1–3^. Despite advances in molecularly targeted agents and hypomethylating therapies, relapse in AML remains common and is largely attributed to leukemic stem and progenitor populations that survive cytotoxic and epigenetic stress through adaptive metabolic programs ^4–6^.

Leukemic stem and progenitor cells (LSCs) exhibit increased reliance on mitochondrial oxidative metabolism, frequently coupled to fatty-acid uptake and β-oxidation (FAO), to sustain stemness-associated programs, quiescence, and redox balance. This contrasts with normal hematopoietic stem cells (HSCs), which are maintained in a predominantly glycolysis-biased, respiration-restricted state ^5,6^. Accordingly, therapy resistance and disease persistence are associated with the selection of oxidative and lipid-dependent metabolic states, including FAO programs supported by the bone marrow microenvironment ^7^.

The clinical relevance of mitochondrial dependence in AML is underscored by the activity of venetoclax, a BCL-2 inhibitor that exploits mitochondrial apoptotic priming and shows substantial efficacy in combination with azacitidine. Metabolic profiling of venetoclax/azacitidine-treated AML demonstrates preferential targeting of stem-cell–enriched compartments and disruption of mitochondrial energy metabolism ^5,8^. However, AML cells display marked metabolic plasticity, enabling the maintenance or re-establishment of mitochondrial oxidative metabolism through alternative lipid acquisition and intracellular lipid trafficking pathways, thereby limiting the durability of metabolic targeting strategies ^9,10^.

Aberrant lipid metabolism is increasingly recognized as a defining feature of AML, encompassing enhanced fatty-acid uptake and functional dependence on FAO to sustain oxidative phosphorylation (OXPHOS) and biosynthetic demands ^6,11^. Beyond fatty acids, emerging evidence from cancer and myeloid leukemia models indicates that sterol metabolism can influence mitochondrial integrity and therapeutic sensitivity ^12,13^. While scavenger receptor–mediated lipid uptake and microenvironmental lipid supply can support FAO-linked resistance programs ^7^, experience with surface-directed targeting in AML highlights that broad antigen expression across normal hematopoietic compartments constrains therapeutic specificity and increases the risk of on-target off-tumor toxicity ^14,15^. In contrast, mitochondrial cholesterol trafficking, mediated by defined endoplasmic reticulum, mitochondria contact sites and capable of modulating mitochondrial bioenergetics and apoptosis sensitivity, represents a mechanistically targetable metabolic axis ^16–19^.

Clever-1 (Stabilin-1) is a multifunctional scavenger receptor expressed on subsets of macrophages and endothelial cells ^20^ and has also been reported in leukemic cells ^21^. Through lipid and apoptotic cargo clearance, Clever-1 contributes to tissue homeostasis but can also promote immunosuppressive programs in cancer ^22^. Because Clever-1 regulates endosomal routing of modified lipoproteins, it may influence intracellular lipid distribution, including pathways relevant to mitochondrial lipid supply. Bexmarilimab, a humanized anti–Clever-1 antibody ^23^, reprograms macrophages toward a pro-inflammatory phenotype ^24^ and shows clinical activity in combination with hypomethylating agents in myeloid malignancies ^25^. Whether Clever-1 also regulates leukemic cell–intrinsic lipid metabolism and mitochondrial fitness remains unknown.

Here, we show that Clever-1 blockade disrupts intracellular lipid trafficking, alters mitochondrial lipid composition, and compromises OXPHOS capacity in AML. By integrating bioenergetic profiling, lipidomics, and proteomics, we identify a previously unrecognized role for Clever-1 in sustaining mitochondrial integrity and metabolic resilience in leukemic cells.

## Materials and methods Cell culture

AML cell lines HL-60, MOLM-13, MV4-11, TF-1 and Kasumi-1 (all from DSMZ) were cultured in RPMI-1640 supplemented with L-glutamine, penicillin–streptomycin (P/S) and fetal bovine serum (FCS) as follows: HL-60, MOLM-13 and MV4-11 in RPMI + 10% FCS; TF-1 in RPMI + 10% FCS + 2 ng/mL GM-CSF; Kasumi-1 in RPMI + 20% FCS. KG-1 (ATCC) cells were grown in IMDM + 20% FCS with L-glutamine and P/S. Cells were maintained at 37 °C, 5% CO□and routinely tested mycoplasma-free.

## Bulk RNA sequencing and gene set enrichment

KG-1, HL-60 and MOLM-13 cells were cultured with bexmarilimab (50 µg/mL, Abzena) or human IgG4 isotype control (BioXcell) for 24–48 h under standard conditions. Cells were washed in PBS and pelleted (300 g, 5 min). Total RNA was extracted using NucleoSpin RNA (Macherey-Nagel). RNA quantity/quality were assessed by NanoDrop and Bioanalyzer. Library preparation and sequencing were performed at Novogene. Differential expression analysis and multiple-testing correction (FDR) were carried out by Novogene. Gene set enrichment analysis (GSEA, Broad Institute) was run on preranked gene lists using Hallmark gene sets (h.all.v2024.1), 1000 permutations and default settings; significant pathways were defined as FDR q < 0.05.

## Airyscan super-resolution microscopy and DiANA analysis

KG-1 cells were blocked in 15% horse serum in PBS and stained with Alexa Fluor 488–conjugated bexmarilimab or IgG4 either on ice or at 37 °C for 30 min. Cells were fixed in 4% paraformaldehyde (PFA) and, for intracellular staining, permeabilized with 0.1% Triton X-100 in 30% horse serum. Cells were subsequently stained for 1 h at 4 °C with Alexa Fluor 647–conjugated 9–11 anti–Clever-1 (InVivo Biotech) or isotype control, or with rabbit anti-SLC25A10 (Proteintech) followed by FITC-conjugated anti-rabbit secondary antibody, together with phalloidin and Hoechst. Cells were spun onto Cell-Tak (Corning)–coated co-verslips and mounted in ProLong Gold (Thermo).

Airyscan super-resolution imaging was performed on a Zeiss LSM 880 microscope equipped with a 63×/1.4 NA oil immersion objective using 405, 488, 561, and 633 nm lasers. Z-stacks were acquired with a step size of 0.16 µm and processed using Zen Black software. Three-dimensional segmentation and colocalization analyses were performed in Fiji using the DiANA plugin. Clever-1–positive and mitochondrial objects were segmented using standard thresholding, with border-touching objects excluded. The proportion of mitochondria containing at least one Clever-1–positive object was calculated on a per-cell basis.

## Immunoelectron microscopy

KG-1 cells were fixed in 4% PFA and 0.1% glutaraldehyde, embedded in LR White and sectioned (∼70 nm). Sections were blocked with 10% FCS in PBS and stained with anti–Clever-1 (clone 4G9; Santa Cruz) followed by 10-nm gold–conjugated goat anti-mouse IgG. Control grids omitted primary antibody or used isotype control. Sections were post-stained with uranyl acetate and lead citrate and imaged on a JEOL JEM-1400 TEM at 80 kV. Gold particles within mitochondria were quantified manually in Fiji.

## Mitochondrial isolation

Mitochondria were isolated from KG-1 cells using anti-TOM22 MicroBeads and a human Mitochondria Isolation Kit (Miltenyi Biotec) according to the manufacturer’s protocol. Briefly, 1 × 10□cells were homogenized in ice-cold lysis buffer with Protease Inhibitor Cocktail (Roche), cleared at 700 g (10 min, 4 °C), and incubated with TOM22 MicroBeads in separation buffer. Magnetically labeled mitochondria were enriched on MACS LS columns and pelleted at 13,000 g (2 min, 4 °C). Pellets were used immediately or resuspended in storage buffer.

## Blue Native PAGE of respiratory complexes

Protein content of mitochondrial pellets was measured by DC Protein Assay (Bio-Rad). For Blue Native PAGE (BN-PAGE), mitochondria were solubilized in NativePAGE sample buffer with digitonin (protein:detergent ratio 1:4), clarified at 20,000 g (30 min, 4 °C) and supplemented with G-250 additive (Thermo). Samples were resolved on 3–12% Bis-Tris NativePAGE gels (Thermo) with cathode additive according to the manufacturer’s two-step protocol. Gels were either stained with GelCode Blue (Thermo) to visualize markers or transferred to PVDF in NuPAGE transfer buffer and processed for immunoblotting with OXPHOS antibody cocktail (Abcam) at 1:500 following with TrueBlot anti-mouse HRP (Rockland). Respiratory complexes and supercomplexes were assigned based on described human BN-PAGE migration patterns ^26^.

## Co-immunoprecipitation and proteomic analysis of Clever-1 complexes

KG-1 cells (6 × 10□per condition) were treated for 24 h with bexmarilimab or IgG4, lysed in Triton X-100–based lysis buffer with protease/phosphatase inhibitors, and cleared at 10,000 g. Lysates were precleared with streptavidin M-280 Dynabeads, then incubated with Dynabeads pre-complexed with biotinylated 9-11 anti–Clever-1 or biotinylated rat isotype control overnight at 4 °C. Beads were extensively washed in lysis buffer and TBS and submitted to the Turku Proteomics Facility for on-bead trypsin digestion, peptide cleanup and LC-MS/MS on a Q Exactive HF mass spectrometer. Spectra were searched with Proteome Discoverer 3.0 software (Thermo) connected to an in-house server running the Mascot 2.8.3 software (Matrix Science) against SwissProt version 2023_01 (Homo sapiens). For each protein, the 9-11 pulldown abundance was divided by isotype abundance to yield a quotient; proteins with quotient < 4 were excluded. Differences in abundance and purity sum between bexmarilimab and control were calculated, and CRAPome scores retrieved to flag common contaminants.

## Western blotting

For validation of interactors and analysis of mitochondrial fractions, proteins were eluted from Dynabeads or mitochondrial pellets in Laemmli sample buffer with DTT, resolved on 4–20% TGX gels (Bio-Rad) and transferred to PVDF using a Trans-Blot Turbo system or wet transfer. Membranes were blocked in 5% milk in TBST and probed with primary antibodies against IMPDH2 (Proteintech), ATAD3A/B (Proteintech), ATP6V0A2 (Invitrogen), Clever-1 (clone 4G9), PDI (clone 1D3; Enzo) or GAPDH (clone 6C5; Hytest). HRP-conjugated secondary antibodies were goat anti-rabbit IgG (DAKO), anti-mouse IgG for IP (Abcam), or mouse anti-rabbit IgG TrueBlot. Signals were detected with ECL (Cytiva). Where necessary, membranes were stripped in glycine/SDS/Tween buffer, reblocked and reprobed.

## Flow cytometry

To characterize Clever-1 expression in AML cell lines, cells were stained with Alexa Fluor 647–conjugated anti-human Clever-1 (9-11; in-house conjugated, 10 µg/mL) to assess surface expression (EC) or total (intracellular) Clever-1 expression after fixation and permeabilization. Clever-1 expression was quantified as median fluorescence intensity following exclusion of dead cells (Fixable Viability Dye eFluor 506, Invitrogen).

For mitochondrial flow cytometry, KG-1 cells were cultured with IgG4 or bexmarilimab (50 µg/mL) for 24 h, labeled for 1 h with MitoView Fixable 640 (Biotium) and processed for mitochondrial MACS isolation. Isolated mitochondria were fixed in 4% PFA and, where indicated, permeabilized with 0.1% Triton X-100. Blocking and staining were performed in 1% BSA/1 mM EDTA PBS. Mitochondria were stained with biotinylated 9-11 anti–Clever-1, anti-hIgG4-FITC and streptavidin-PE. Mitochondria were identified based on forward/side scatter characteristics and MitoView Fixable 640 positivity.

For acLDL uptake, KG-1 cells were pretreated with IgG4 or bexmarilimab for 24 h, pulsed with A488-acLDL for 2 h and MitoView for the final hour. For whole-cell measurements, cells were washed, fixed in 1% PFA and treated with 0.2% trypan blue to quench surface fluorescence. For mitochondrial acLDL analysis, mitochondria were isolated and fixed in 1% PFA. AcLDL uptake was quantified as percentage of acLDL-positive events and geometric mean fluorescence intensity. All samples were acquired on BD FACSFortessa or FACSymphony instruments. Data were analyzed in FlowJo software v10.10.0 (BD).

## Mitochondrial lipidomics

KG-1 cells were cultured with IgG4 or bexmarilimab (50 µg/mL, 24 h), and mitochondria were isolated as above. Pellets were resuspended in 0.9% NaCl; a small aliquot was kept for normalization. Lipids were extracted in CHCl□:MeOH (2:1) and analyzed by UHPLC-MS/MS using an ACQUITY UPLC BEH C18 column on a 6600 UHPLC (Sciex) coupled to a Q Exactive HF mass spectrometer. A binary gradient of aqueous and organic mobile phases containing ammonium acetate and formic acid was applied, followed by re-equilibration between runs. Lipid standards were used for peak identification, and experiments were performed in triplicate.

## Oxygen consumption rate (OCR) assessment

AML cells were treated with bexmarilimab or IgG4 for 48 h, washed and resuspended in Seahorse XF RPMI medium (pH 7.4) containing 10 mM glucose, 2 mM L-glutamine and 1 mM sodium pyruvate. Cells (7.5 × 10□/well) were seeded onto Cell-Tak–coated XF96 plates and allowed to attach and equilibrate at 37 °C in a CO□-free incubator for 1 h, following the manufacturer’s protocol. OCR was measured using the Seahorse XF Mito Stress Test (Ag-ilent) with sequential injections of oligomycin, FCCP (co-injected with additional pyruvate) and rotenone/antimycin A. Final inhibitor concentrations were optimized per line. Hoechst was added at the end of the run, and OCR values were normalized to nuclei counts by automated imaging.

## Transmission electron microscopy

HL-60, MOLM-13 and KG-1 cells were cultured for 48 h with IgG4 or bexmarilimab, washed in PBS and pelleted in BEEM capsules. Pellets were fixed in 2.5% glutaraldehyde, dehydrated through graded ethanol and embedded in epoxy resin. Ultrathin sections (∼70 nm) were cut on a Leica UC7 ultramicrotome, mounted on EM grids and post-stained with uranyl acetate and lead citrate. Images were acquired on a JEOL JEM-1400 TEM at 80 kV. Mitochondria were segmented using Microscopy Image Browser (MiB; MATLAB version 2.91) ^27^ with the Segment Anything plugin (SAM2, version 2.1; Meta AI), and mitochondrial area and filled area were quantified for each field.

## Flow cytometric analysis of mitochondrial dysfunction

KG-1 cells were cultured with IgG4 or bexmarilimab (50 µg/mL, 48 h) in (i) complete IMDM + 20% FCS, (ii) minimal RPMI without glucose/glutamine + 20% FCS, (iii) minimal RPMI + 20% human AB serum, or (iv) minimal RPMI + 20% delipidized human serum (SP1010, Veritas Innovation). Cells were stained with Fixable Viability Dye eFluor 450, then with MitoTracker Green FM and MitoTracker Red CMXRos in pre-warmed medium. After washing, cells were analyzed on a BD FACSFortessa. Functional mitochondria were defined as MitoTracker Green^high/MitoTracker Red^high and dysfunctional mitochondria as MitoTracker Green^high/MitoTracker Red^low in FlowJo as described previously ^28^.

## Statistical analysis

Unless otherwise stated, statistical analyses were performed in GraphPad Prism (version 10.6.1). Comparisons between two groups used Welch’s t-test; analyses with more than two groups used one-way or two-way ANOVA with appropriate post hoc tests, as detailed in figure legends. Flow cytometry data are presented as median fluorescence intensity (MFI). Imaging-based measurements (Airyscan/DiANA and TEM–MiB) were treated as independent biological replicates at the image/field level. RNA-seq differential expression and FDR correction were performed by Novogene, and GSEA used Hallmark gene sets from MSigDB. Statistical significance was defined as p < 0.05.

## Results

### Clever-1 blockade reprograms mitochondrial transcriptional programs in AML cells

To characterize the subcellular distribution of Clever-1 in AML, we first examined surface and total protein levels across five AML cell lines by flow cytometry. Expression varied markedly between models, with KG-1 showing the highest abundance (**Fig. 1A**). Given that Clever-1 mediates efficient uptake of bexmarilimab in macrophages, we next investigated whether AML cells likewise support binding and internalization of this clinically developed antibody ^29^. At 4 °C, where endocytosis is halted, immunofluorescence staining revealed bexmarilimab bound to surface Clever-1 restricted to discrete membrane regions. Shifting the cells to 37 °C for 30 min resulted in redistribution of the Clever-1–bexmarilimab complexes into intracellular compartments, consistent with efficient receptor-mediated internalization (**Fig. 1B**). We then investigated how Clever-1 inhibition with bexmarilimab reshapes leukemic transcriptional programs. Bulk RNA-seq was performed on KG-1 cells treated with bexmarilimab for 24 and 48 h. In KG-1 cells, bexmarilimab triggered a pronounced early response at 24 h, with 41 differentially expressed genes (padj < 0.05). Many of the most strongly upregulated transcripts encoded mitochondrial proteins involved in import, translation, and respiratory chain function, including *TOMM7*, *MRPL12*, *HSPE1*, *NDUFB2*, *COX6A1*, and *ATP5IF1* (**Fig. 1C**). Pathway analysis at 24 h showed enrichment of oxidative phosphorylation, hypoxia, and adipogenesis signatures (**Fig. 1C**). By 48 h, only a small number of transcripts remained significantly altered, yet gene-set enrichment demonstrated downregulation of oxidative phosphorylation, fatty acid metabolism, and reactive oxygen species pathways (**Fig. 1D**), suggesting that an early mitochondrial activation phase transitions into a compensatory or stressed metabolic state.

**Figure 1.**
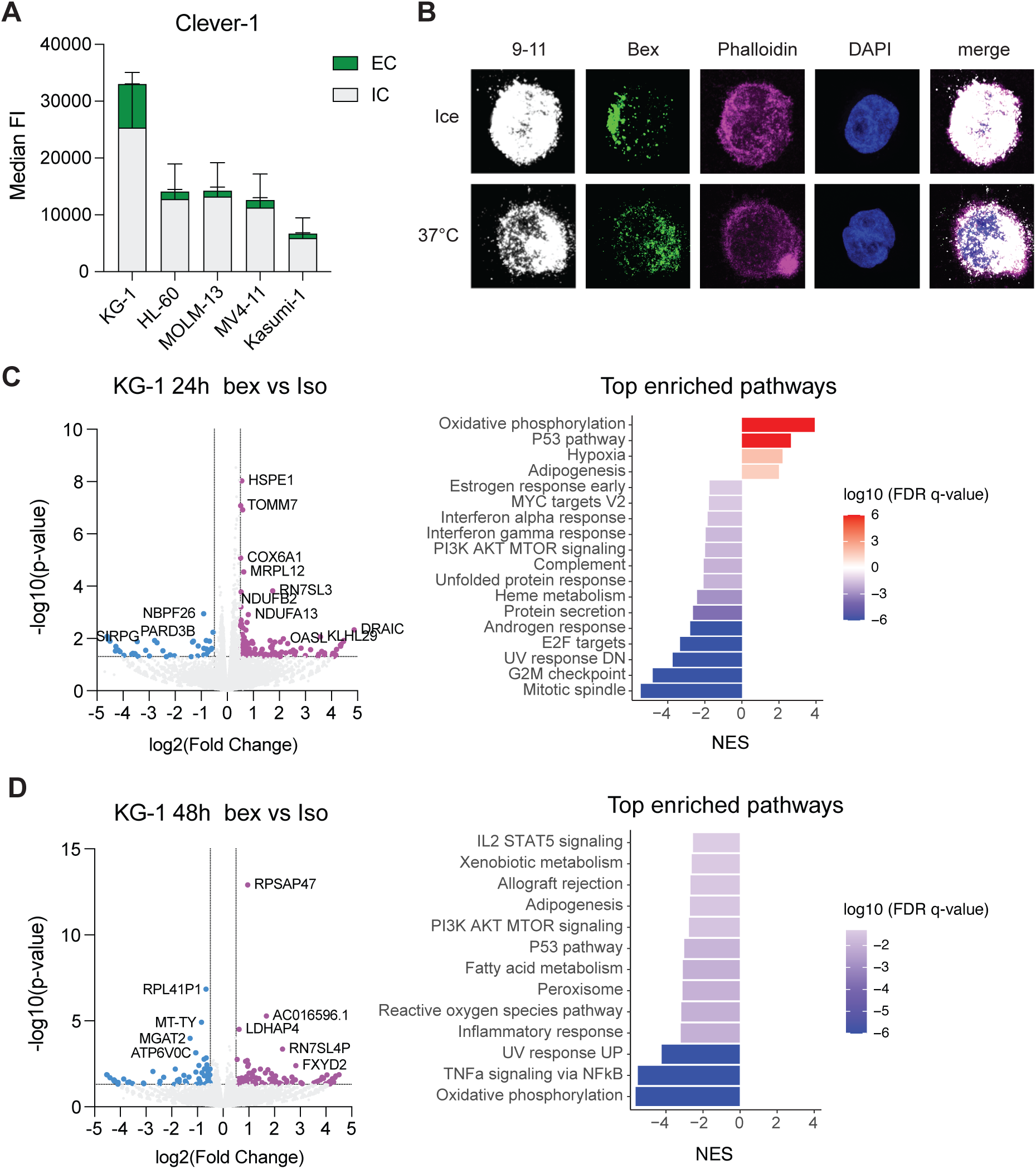
Clever-1 blockade induces transcriptional programs linked to mitochondrial metabolism in AML cells. A, Flow cytometry analysis of surface (EC) and intracellular (IC) Clever-1 expression in AML cell lines. (n=15) **B,** Airyscan super-resolution microscopy images of KG-1 cells showing (bex, green) internalization after 30 min incubation at 37°C or 4°C. Cells were fixed, permeabilized, and stained with anti–Clever-1 (9-11, white), phalloidin (F-actin, magenta), and DAPI (nuclei, blue). **C**–**D,** Volcano plots and pathway analyses of differential gene expression in KG-1 cells treated with bexmarilimab versus isotype (iso) control antibody for 24 h (C) and 48 h (D) (n=3). Upregulated and downregulated genes are highlighted in magenta and blue, respectively. Differential expression was determined by RNA-seq with Benjamini–Hochberg false discovery rate (FDR) correction; genes with adjusted p values (FDR) < 0.05 were considered significant in C. Gene set enrichment analysis (GSEA) was performed on preranked gene lists using Hallmark gene sets with 1,000 permutations; pathways were considered significant at nominal p < 0.05 in D.

## Clever-1 localizes to mitochondria and associates with mitochondrial proteins

Building on the transcriptional evidence for mitochondrial perturbation after Clever-1 blockade, we therefore asked whether the receptor localizes in proximity to mitochondria. Airyscan super-resolution imaging using the inner-membrane carrier SLC25A10 identified discrete regions where Clever-1–positive puncta overlapped with mitochondrial structures in KG-1 cells (**Fig. 2A**). Quantitative 3D segmentation and proximity analysis using the DiANA algorithm showed that roughly one-third of mitochondria (∼35%) contained overlapping or contacting Clever-1 signal, significantly higher than in isotype-treated controls (∼2%) (**Fig. 2B**). Moreover, we performed immunoelectron microscopy at ultrastructural resolution using a primary antibody against Clever-1 and gold-conjugated secondary antibodies for detection. Clever-1 labeling was detected in association with mitochondrial structures, and quantitative analysis revealed increased mitochondrial localization compared with control antibody staining (**Fig. 2C**). These complementary imaging approaches support that Clever-1 localizes to mitochondria in AML cells.

**Figure 2.**
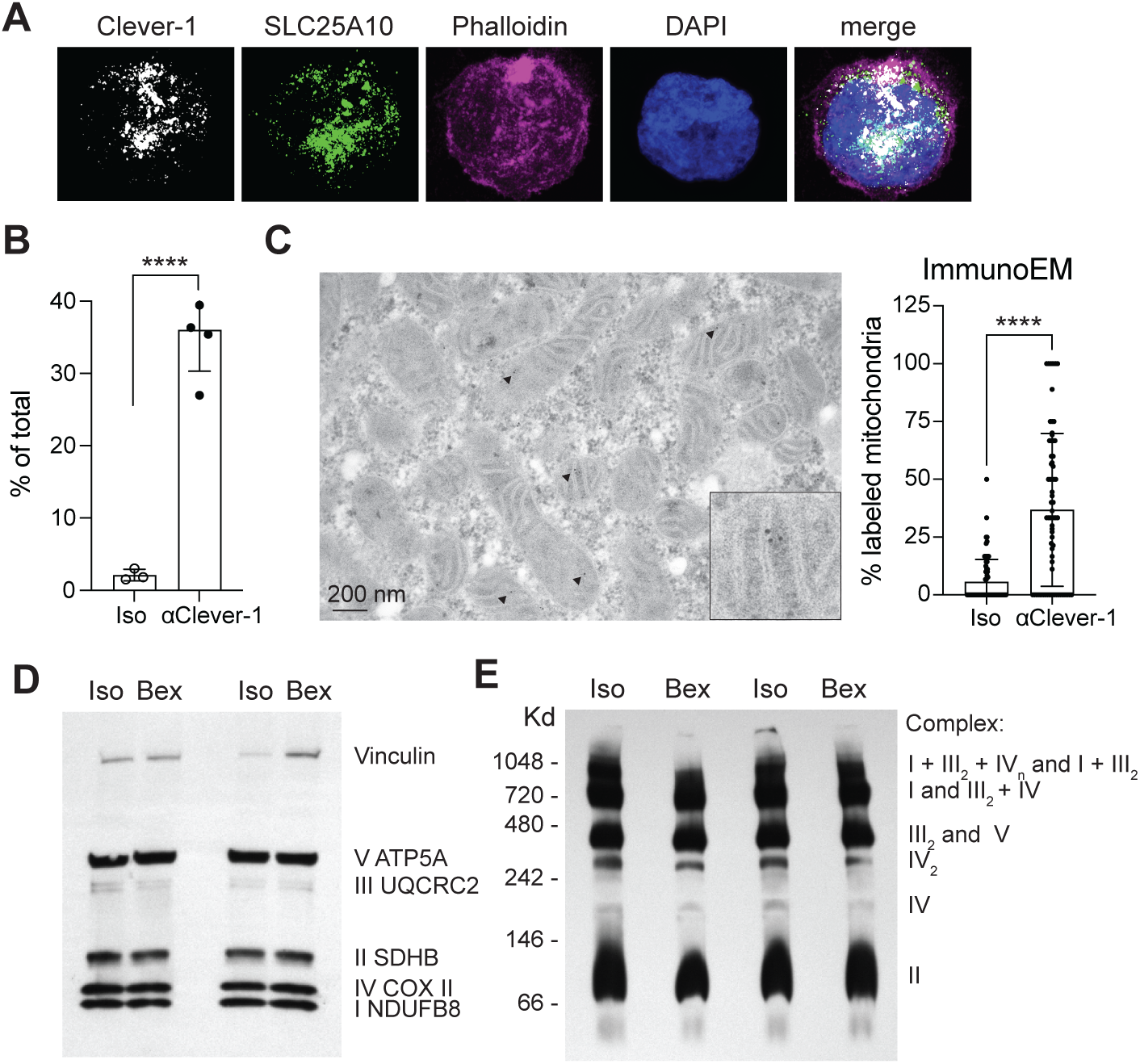
Mitochondrial Clever-1 influences respiratory complex IV assembly in AML cells. A, Airyscan super-resolution images of KG-1 leukemia cells showing Clever-1 (white) in partial spatial overlap with the inner mitochondrial membrane marker SLC25A10 (green). **B,** Quantification of Clever-1 mitochondrial association using 3D segmentation and colocalization analysis (DiANA), showing the percentage of mitochondria containing ≥1 Clev-er-1–positive punctum. **C,** Transmission electron microscopy with immunogold labeling demonstrating mitochondrial localization of Clever-1 in KG-1 cells, quantified as the percentage of gold-labeled mitochondria across 59 independently acquired TEM images. Arrowheads indicate immunogold particles in mitochondria. **D,** Western blot analysis of enriched mitochondria showing steady-state levels of respiratory chain complex subunit proteins (Complexes I–V) after 24 h bexmarilimab treatment. Vinculin served as a loading control. **E,** Blue Native PAGE analysis demonstrating altered assembly of respiratory chain complexes and supercomplexes following 24 h bexmarilimab treatment, with a consistent reduction in Complex IV and IV□ assembly. *P* **** < 0.0001.

On the basis of the Airyscan super-resolution and electron microscopy evidence for mitochondrial localization of Clever-1 in KG-1 cells, we next examined whether bexmarilimab treatment alters the organization of the mitochondrial respiratory chain. Using conventional SDS-PAGE, we observed no detectable changes in the abundance of individual electron transport chain proteins across duplicate experiments in enriched mitochondria. In contrast, Blue Native PAGE, which preserves intact respiratory chain complexes, revealed a subtle but reproducible reduction in the assembly of Complex IV and its dimer (IV ) following bexmarilimab treatment (**Fig. 2D–E**).

### Bexmarilimab alters the Clever-1 interactome and affects mitochondrial-associated protein complexes

Co-immunoprecipitation of Clever-1 with the 9-11 antibody followed by mass spectrometry identified several proteins whose association with Clever-1 was altered by bexmarilimab treatment (**Fig. 3A**, Supplementary data file). Validation by repeated co-immunoprecipitation and Western blot analysis confirmed that Clever-1, ATAD3, and IMPDH2 consistently showed reduced precipitation with the 9-11 antibody after bexmarilimab treatment (**Fig. 3B**). To further evaluate these interactions, the pulldown was inverted by using biotinylated antibodies against ATAD3 or IMPDH2 followed by Western blotting for Clever-1. In contrast to the 9-11 pulldown, both ATAD3 and IMPDH2 precipitated increased amounts of Clever-1 from bexmarilimab-treated KG-1 cells compared with isotype controls, with ATAD3 showing the most pronounced effect (**Fig. 3C**). This opposing pattern suggests that bexmarilimab alters the interaction between Clever-1 and these proteins, potentially by enhancing their association while partially masking the 9–11 antibody epitope.

**Figure 3.**
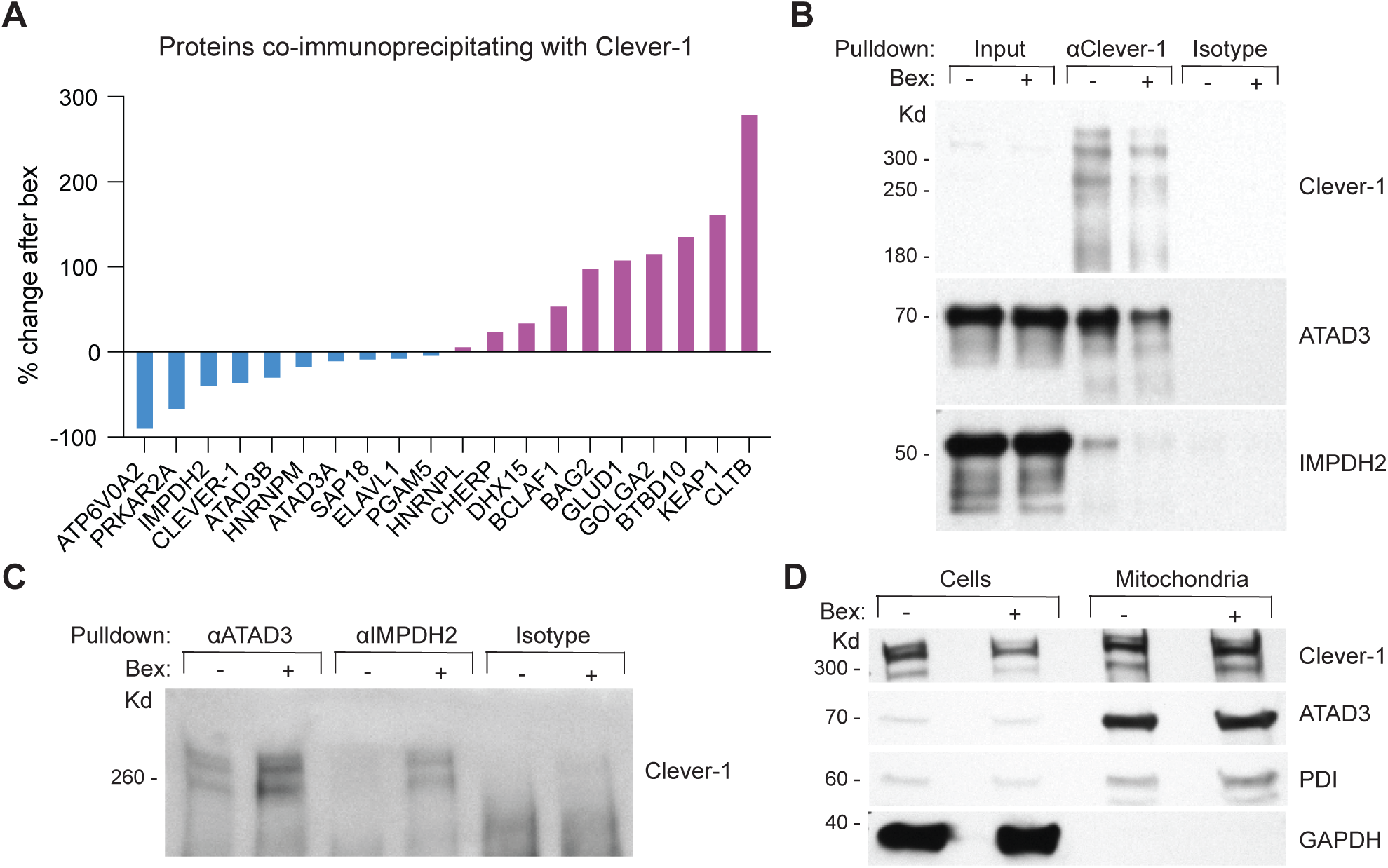
Bexmarilimab alters the interaction between Clever-1 and mitochondrial-associated proteins. A, Anti–Clever-1 (9–11) pulldown and mass spectrometry analysis of proteins associating with Clever-1 in KG-1 cells after 24 h bexmarilimab treatment. **B,** Western blot analysis of ATAD3 and IMPDH2 following anti–Clever-1 immunoprecipitation from bexmarilimab-treated KG-1 cells. Input lysate and isotype IgG pulldown serve as controls. **C,** Western blot detection of Clever-1 following anti-ATAD3 and anti-IMPDH2 immunoprecipitation from bexmarilimab-treated KG-1 cells. Isotype IgG pulldown serves as control. **D,** Western blot of ATAD3 and Clever-1 in whole-cell lysates and mitochondrial fractions. PDI (endoplasmic reticulum) and GAPDH (cytosolic) were used as compartmental markers to assess fraction purity. The Western blot images have been cropped and the original ones can be found in Supplemental Fig 1.

Given that ATAD3 is a mitochondrial membrane protein and Clever-1 shows prominent mitochondrial localization (**Fig. 3A–B**), we subsequently examined whether bexmarilimab affects the distribution of these proteins within mitochondria. Mitochondria isolated from bexmarilimab- and isotype-treated KG-1 cells were analyzed by Western blot. While bexmarilimab reduced total Clever-1 levels in whole-cell lysates, mitochondrial Clever-1 abundance remained unchanged, and mitochondrial ATAD3 levels were similarly unaffected (**Fig. 3D**). The mitochondrial preparations showed minimal contamination by other organelles, and bexmarilimab was undetectable in the mitochondrial fraction (data not shown). These findings indicate that bexmarilimab preferentially reduces non-mitochondrial Clever-1, consistent with internalization and degradation of bexmarilimab–Clever-1 complexes while preserving the mitochondrial-associated Clever-1 pool.

### Clever-1 inhibition reduces acLDL uptake and disrupts lipid trafficking to mitochondria

Because Clever-1 and ATAD3 are both linked to cholesterol transport and localize to mitochondria, we investigated whether Clever-1 blockade affects the delivery of lipid cargo to mitochondria. KG-1 cells were pretreated with bexmarilimab, pulsed with fluorescent acLDL, and analyzed by flow cytometry at the whole-cell level and in enriched mitochondria. Bexmarilimab reduced cellular acLDL uptake compared with isotype control (**Fig. 4A**) and caused an even more pronounced decrease in acLDL signal in the mitochondrial fraction, indicating impaired delivery of lipoprotein-derived lipids to mitochondria.

**Figure 4.**
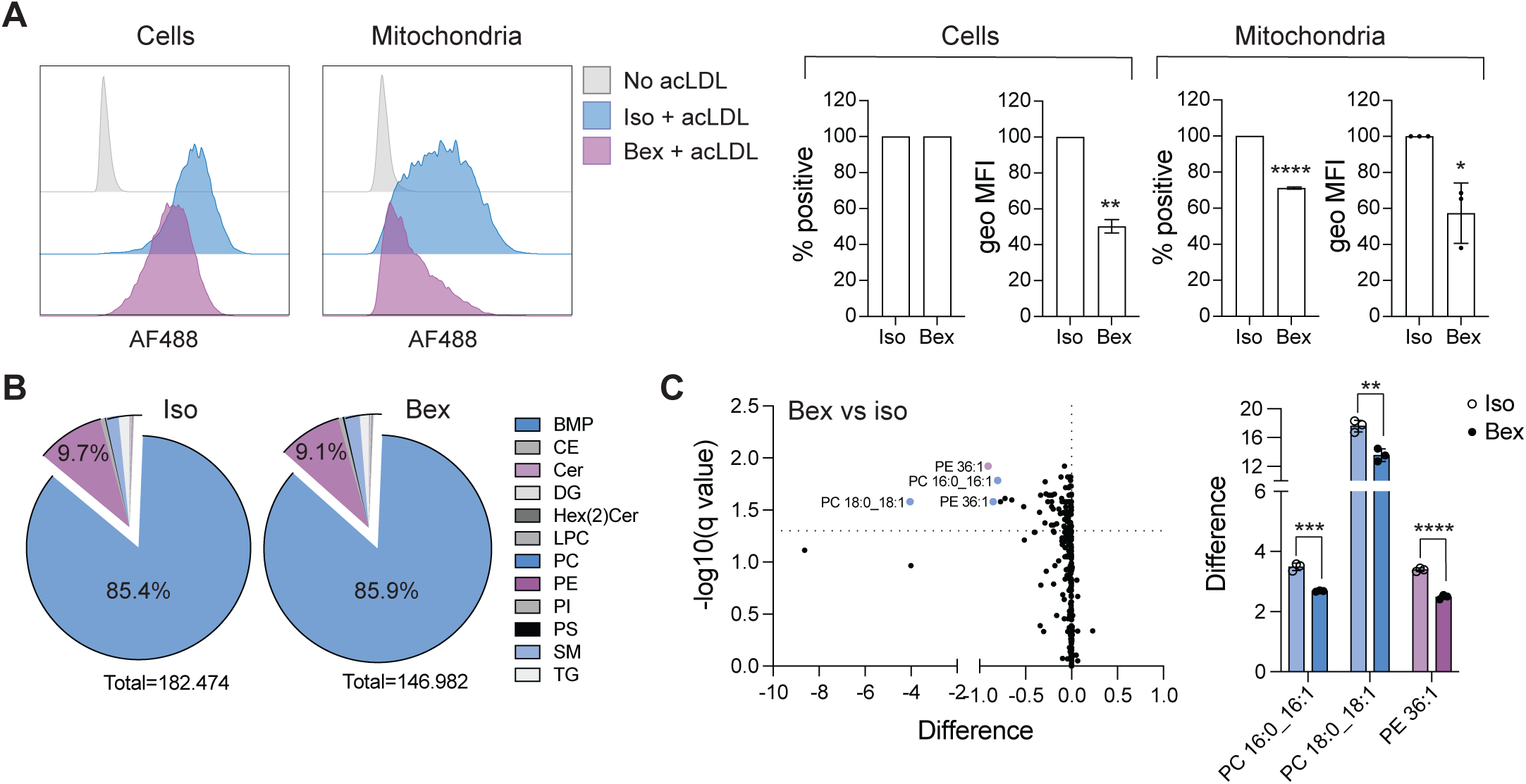
Bexmarilimab reduces lipid trafficking to mitochondria in AML cells.A, Flow cytometry analysis and quantification of acetylated (ac)LDL uptake in whole KG-1 cells and in mitochondria-enriched fractions following isotype or bexmarilimab treatment. Mitochondria were identified by forward/side scatter characteristics and MitoView Fixable 640 positivity prior to analysis of acLDL fluorescence. Shown are representative histograms and quantification of the percentage of acLDL-positive events and geometric mean fluorescence intensity (geo MFI). Data were normalized to the isotype condition within each experiment and analyzed using a one-sample Student’s *t*-test. **B,** Targeted lipidomics analysis of mitochondria-enriched fractions isolated from KG-1 cells following isotype or bexmarilimab treatment. Pie charts summarize the relative proportions of mitochondrial lipid classes, with phosphatidylcholines (PC, blue) and phosphatidylethanolamines (PE, purple) representing the dominant species. Other lipid classes identified include: BMP = bis(monoacylglycero)phosphates, CE = cholesteryl esters, Cer = ceramides, DG = diacylglycerols, Hex(2)Cer = dihexosylceramides, LPC = lysophosphatidylcholines, PI = phosphatidylinositols, PS = phosphatidylserines, SM = sphingomyelins, TG = triglycerides. **C,** Differential abundance analysis of mitochondrial lipid species following bexmarilimab treatment, highlighting selected significantly reduced phosphatidylcholine (PC) and phosphatidylethanolamine (PE) species. Statistical significance was assessed using multiple unpaired *t*-tests. *P* ** < 0.01, *** < 0.001, **** < 0.0001.

To assess downstream consequences for mitochondrial lipid composition, we performed targeted lipidomic profiling of mitochondria isolated from bexmarilimab-treated KG-1 cells. Bexmarilimab induced subtle but significant reductions in selected mitochondrial lipid species (**Fig. 4B–C**, Supplementary Fig. 2). Notably, bexmarilimab did not alter cell-surface Clever-1 expression, strongly inhibit acLDL binding to Clever-1, or change overall mitochondrial mass as assessed by MitoView 640 staining (Supplementary Fig. 3). Together, these data support a model in which bexmarilimab partially impairs Clever-1–mediated trafficking of lipid cargo to mitochondria rather than globally depleting mitochondria or abolishing acLDL binding.

### AML cell lines with high baseline OXPHOS are most sensitive to mitochondrial respiratory suppression by bexmarilimab

To determine how Clever-1 inhibition affects mitochondrial respiration across genetically and phenotypically diverse AML cell lines, we performed a Seahorse XF mitochondrial stress test after 48 h exposure to bexmarilimab under standard cell culture conditions. The AML panel exhibited substantial heterogeneity in baseline mitochondrial respiratory activity and spare respiratory capacity, and these metabolic profiles did not align with French–American–British (FAB) classification or canonical mutational annotations, nor the amount of Clever-1 on the cell surface (**Fig. 5A**; **Fig. 1A**). Cell lines such as HL-60 and KG-1 displayed relatively high mitochondrial respiratory activity under control conditions, whereas Kasumi-1 and MV4-11 demonstrated more restrained respiratory flux, consistent with prior reports demonstrating substantial inter-cell-line heterogeneity in mitochondrial respiratory capacity and spare respiratory reserve among AML models ^30^.

**Figure 5.**
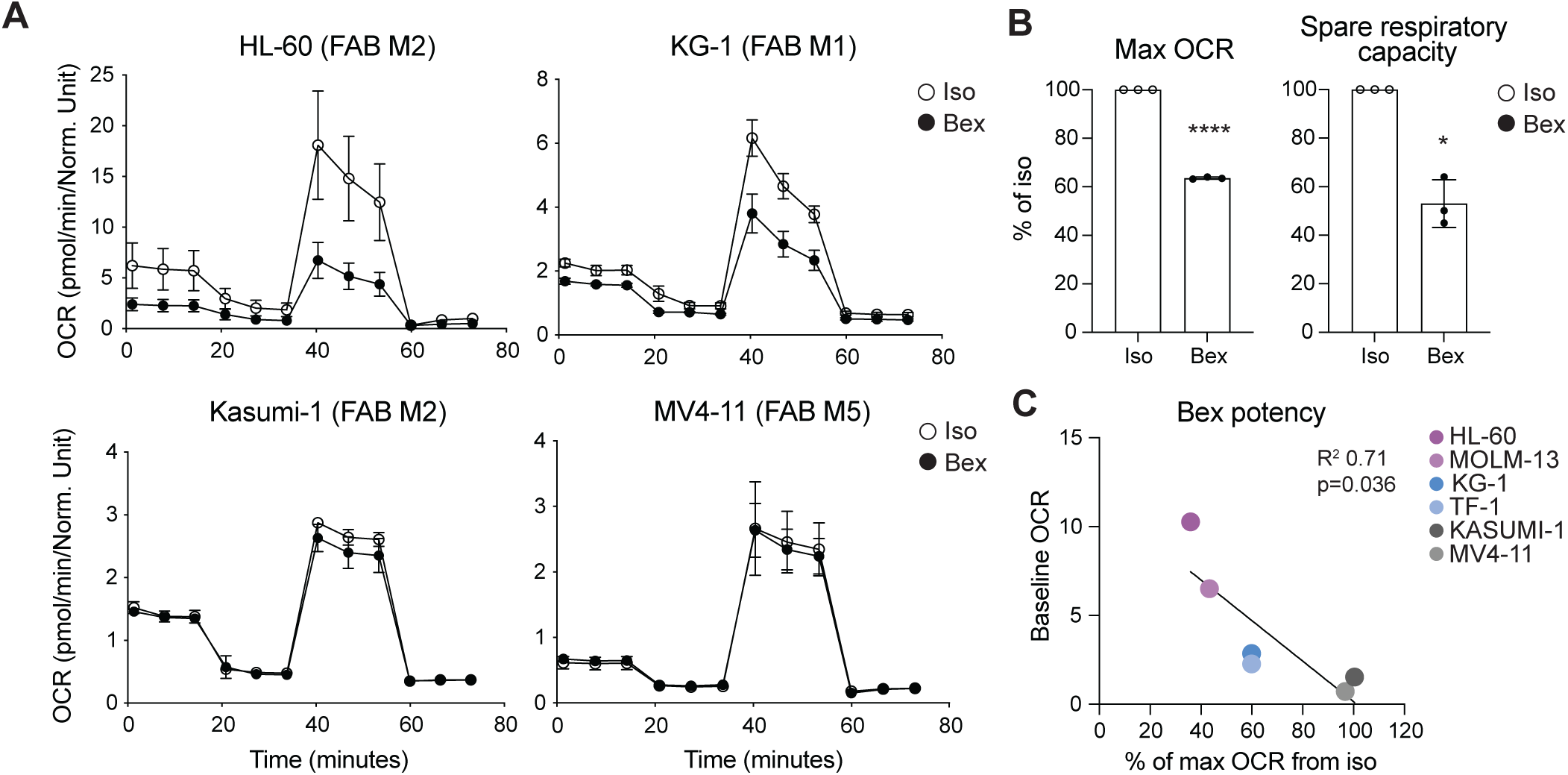
Bexmarilimab impairs mitochondrial respiration in AML cell lines with high baseline oxidative phosphorylation. A, Seahorse XF Mitochondrial Stress Test analysis of oxygen consumption rate (OCR) in AML cell lines (HL-60, KG-1, Kasumi-1, and MV4-11) cultured under standard conditions and treated with isotype control or bexmarilimab for 48 h prior to assay. Sequential injections of oligomycin (ATP-linked respiration; ∼20 min), FCCP (maximal respiration; ∼35 min), and rotenone/antimycin A (non-mitochondrial respiration; ∼60 min) are indicated. Traces reflect mean ± SEM. OCR values were normalized to cell number determined by Hoechst nuclear staining at the end of the assay. The French–American–British (FAB) classification is annotated for each cell line. **B,** Quantification of maximal OCR and spare respiratory capacity in KG-1 cells across three independent experiments. Data are normalized to isotype control. **C,** Relationship between each cell line’s baseline OCR and the magnitude of bexmarilimab-mediated reduction in maximal respiration. Baseline OCR represents the pre-injection measurement averaged across five replicate wells per condition within a single experiment, demonstrating preferential sensitivity in AML cell lines with high baseline OXPHOS activity. *P* * < 0.05, *** < 0.001.

Bexmarilimab treatment reduced oxygen consumption rate (OCR) across multiple AML cell lines, albeit with variable magnitude. KG-1 cells exhibited consistent suppression, with reductions observed in basal OCR, FCCP-stimulated maximal respiration, and spare respiratory capacity (**Fig. 5B**). Other cell lines with high baseline OXPHOS activity showed qualitatively similar, though more variable, decreases in mitochondrial respiration. Notably, comparison of baseline mitochondrial respiration activity with the extent of bexmarilimab-induced suppression revealed a significant inverse correlation (R² = 0.71, p = 0.036), indicating that cell lines with higher basal mitochondrial activity were more sensitive to inhibition of maximal OCR following Clever-1 blockade (**Fig. 5C**). Collectively, these data indicate that bexmarilimab can modulate mitochondrial respiration in AML cells, with more pronounced effects in cell lines characterized by high baseline OXPHOS activity.

## Clever-1 inhibition disrupts mitochondrial ultrastructure and increases dysfunctional mitochondria under metabolic stress

To characterize how Clever-1 inhibition alters mitochondrial morphology in AML cells, we examined HL-60, MOLM-13 and KG-1 cells cultured under standard conditions and treated with bexmarilimab for 48 h, followed by transmission electron microscopy using conventional glutaraldehyde fixation, ultrathin sectioning, and contrast staining. In isotype-treated cells, mitochondria appeared elongated with dense matrices and intact cristae. In contrast, bexmarilimab-treated HL-60 cells exhibited ultrastructural alterations, including increased mitochondrial heterogeneity characterized by a higher abundance of small, rounded mitochondria together with reduced matrix electron density and disrupted cristae architecture (**Fig. 6A**). Notably, concentric “onion-like” cristae, characterized by multilamellar cristae organization, were detected exclusively in the bexmarilimab condition, accompanied by an increased abundance of vacuolar structures containing double-membrane mitochondrial remnants, suggesting accumulation of structurally abnormal mitochondria.

**Figure 6.**
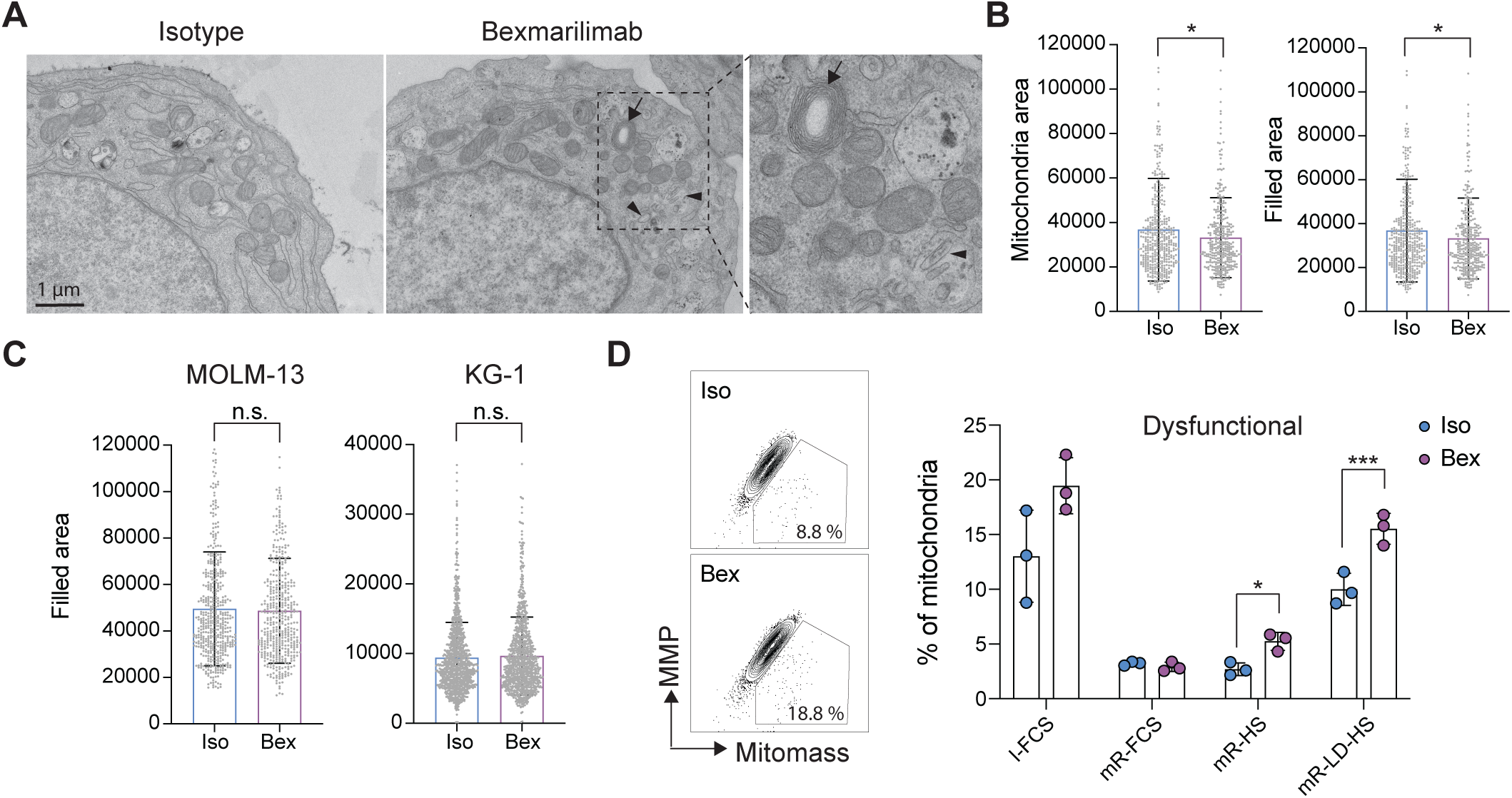
Metabolic stress unmasks bexmarilimab-induced mitochondrial dysfunction. **A**, Representative transmission electron microscopy (TEM) images of HL-60 cells treated with isotype control or bexmarilimab for 48 h under standard culture conditions. Cells were fixed in glutaraldehyde, ultrathin-sectioned (∼70 nm), and contrast-stained with uranyl acetate and lead citrate. Bexmarilimab-treated cells exhibit increased mitochondrial heterogeneity, including small, rounded mitochondria with reduced matrix electron density and disrupted cristae architecture. Concentric onion-like cristae are indicated by arrows, and vacuolar structures containing internal membranous material consistent with mitochondrial remnants are indicated by arrowheads. **B,** Quantification of mitochondrial area and filled (electron-dense) area in HL-60 cells based on semi-automated segmentation of TEM images using MiB in combination with the Segment Anything algorithm. Welch’s t-test. **C,** Parallel quantification of filled mitochondrial area in MOLM-13 and KG-1 cells treated with isotype or bexmarilimab for 48 h, showing no significant ultrastructural alterations under standard culture conditions. **D,** Flow cytometric assessment of mitochondrial membrane potential (MMP) and mitochondrial mass in KG-1 cells using potential-dependent (MitoTracker Red) and po-tential-independent (MitoTracker Green) dyes. Representative plots and quantification of dysfunctional mitochondria (MitoTracker Green^high / MitoTracker Red^low) in cells cultured in complete medium (IMDM + 20% FCS), minimal RPMI lacking glucose and gluta-mine supplemented with FCS or human serum (HS), or minimal RPMI with 20% lipid-depleted HS. Statistical significance determined by Welch’s t-test. *P* * < 0.05, *** < 0.001.

To quantify these morphological alterations, we performed semi-automated analysis of mitochondrial morphology on TEM micrographs using MiB in combination with the Segment Anything algorithm. In HL-60 cells, this analysis revealed a significant reduction in mitochondrial area, reflecting a shift toward smaller mitochondrial profiles, along with decreased filled (electron-dense) area, following bexmarilimab treatment (**Fig. 6B**). These quantitative changes are consistent with altered mitochondrial structure, including cristae disorganization and loss of internal density. In contrast, parallel quantitative analyses in MOLM-13 and KG-1 cells did not reveal significant mitochondrial alterations under standard culture conditions (**Fig. 6C**).

Given the weaker ultrastructural responses in MOLM-13 and KG-1 cells, we then considered whether mitochondrial stress induced by Clever-1 inhibition might become more apparent when cells are forced to rely more heavily on mitochondrial lipid metabolism. Based on our findings that bexmarilimab impairs lipid trafficking to mitochondria, we asked whether this metabolic shift would enhance sensitivity to Clever-1 inhibition. We therefore evaluated mitochondrial function in KG-1 cells across different metabolic conditions.

Using flow-cytometric assessment of mitochondrial mass and membrane potential with potential-independent (MitoTracker Green) and potential-dependent (MitoTracker Red) probes, we quantified the proportion of mitochondria with reduced membrane potential, a marker of mitochondrial dysfunction, under each condition. Bexmarilimab modestly increased mitochondrial dysfunction in complete medium; however, the effect became substantially more pronounced under nutrient-restrictive conditions (minimal RPMI lacking glucose and glutamine) and was further exacerbated when lipid-depleted human serum was used (**Fig. 6D**). These findings indicate that Clever-1 inhibition compromises mitochondrial stability and function, particularly in AML cells under metabolic conditions that increase reliance on endogenous lipid metabolism.

Together, the ultrastructural and functional analyses indicate that Clever-1 plays a role in lipid and cholesterol handling required to maintain mitochondrial integrity, and that inhibition of Clever-1 with bexmarilimab destabilizes mitochondrial architecture, promotes accumulation of dysfunctional mitochondria, and sensitizes AML cells to metabolic stress.

## Discussion

This study identifies Clever-1 as a previously unrecognized regulator of mitochondrial integrity and metabolic resilience in AML. By integrating transcriptional profiling, high-resolution imaging, proteomics, lipidomics, and functional bioenergetic analyses, we demonstrate that Clever-1 blockade disrupts intracellular lipid routing, alters mitochondrial membrane composition and ultrastructure, and selectively compromises OXPHOS in metabolically oxidative AML states. Together, these findings position Clever-1 at the interface of lipid handling and mitochondrial fitness in leukemic cells.

Clever-1 is best characterized as a scavenger receptor regulating immunosuppressive myeloid programs ^20^. However, high Clever-1 mRNA expression in AML is associated with poor survival and disease recurrence ^31^, while protein-level analyses show expression in progenitor-enriched myeloid leukemia populations without defining a leukemic cell–intrinsic role ^21^. Our data extend these observations by demonstrating mitochondrial localization of Clever-1 in AML cells and revealing direct metabolic effects independent of its established immunomodulatory functions. The mitochondrial localization was observed under basal conditions and was not restricted to antibody-treated cells, indicating that mitochondrial association is an intrinsic feature of Clever-1 biology in AML rather than an artifact of receptor engagement. These findings challenge the prevailing view of Clever-1 as primarily an endosomal scavenger receptor with dynamic membrane recycling ^32^ and suggest additional intracellular roles relevant to leukemic cell metabolism.

Proteomic interrogation of the Clever-1 interactome identified altered association with ATAD3 and IMPDH2 following bexmarilimab treatment. ATAD3 is of particular interest given its established roles at mitochondrial membranes and ER–mitochondria contact sites, where it participates in cholesterol trafficking and sterol channeling ^33–36^. Genetic and in vivo studies link ATAD3 to inner mitochondrial membrane organization, cristae structure, and mitochondrial cholesterol handling ^37–40^. Although these functions have not been examined specifically in AML, the association between Clever-1 and ATAD3, together with preserved mitochondrial ATAD3 abundance but altered mitochondrial structure and lipid composition following Clever-1 blockade, is consistent with a model in which Clever-1 supports lipid delivery pathways converging on ATAD3-regulated membrane organization.

IMPDH2, a rate-limiting enzyme in guanine nucleotide synthesis, was also identified as a Clever-1–associated protein. Beyond nucleotide metabolism, IMPDH2 has been implicated in proliferative signaling, apoptosis regulation, and therapy resistance in cancer, including hematologic malignancies ^41–44^. While IMPDH2 is not a mitochondrial enzyme, regulation of guanine nucleotide pools can influence GTP-dependent signaling and trafficking processes intersecting with mitochondrial dynamics and stress responses, suggesting that altered Clev-er-1–IMPDH2 association may reflect broader metabolic rewiring rather than a direct mitochondrial function.

Functionally, Clever-1 blockade reduced uptake of modified LDL and, more strikingly, diminished trafficking of lipid cargo to mitochondria. Lipidomic profiling revealed selective depletion of mitochondrial lipid species after Clever-1 blockade, without evidence of reduced mitochondrial content. Mitochondrial lipid supply is a regulated process coordinated at membrane contact sites, particularly ER–mitochondria interfaces, and involves multiple trafficking proteins ^45–48^. While we did not directly interrogate contact-site machinery, the association of Clever-1 with ATAD3 raises the possibility that Clever-1–dependent trafficking intersects with contact-site–mediated lipid delivery rather than acting solely through endolysosomal routing.

Consistent with impaired lipid delivery, Clever-1 blockade induced pronounced mitochondrial ultrastructural abnormalities, including cristae rarefaction, reduced matrix density, and accumulation of small, rounded mitochondria, accompanied by increased mitophagic structures and a higher fraction of depolarized mitochondria under metabolic stress. At the level of the respiratory chain, blue native PAGE revealed a subtle but consistent reduction in assembly of Complex IV and IV without changes in total OXPHOS protein abundance. Complex IV assembly is increasingly recognized as a regulated and AML-relevant vulnerability ^49^, and its stability depends on the lipid environment of the inner mitochondrial membrane, providing a mechanistic link between impaired lipid delivery, cristae disorganization, and reduced respiratory competence.

Transcriptional and functional metabolic analyses together indicate a dynamic but ultimately maladaptive mitochondrial response to Clever-1 blockade. RNA sequencing revealed early induction of mitochondrial import, translation, and OXPHOS-associated transcripts, consistent with a compensatory attempt to preserve mitochondrial function, followed by suppression of oxidative phosphorylation and fatty-acid metabolism pathways at later time points. Seahorse analyses mirrored this trajectory, demonstrating reduced basal and maximal respiration and diminished spare respiratory capacity. Importantly, the magnitude of respiratory suppression correlated inversely with baseline OXPHOS activity, indicating that OXPHOS-high AML cell lines are particularly dependent on Clever-1–supported mitochondrial metabolism. Notably, AML cell lines with relatively high mitochondrial respiratory capacity, including HL-60 and KG-1, have previously been reported to differ in mitochondrial ultrastructure, ER–mitochondria contact organization, and sensitivity to mitochondrial perturbation, features that have been linked to differentiation state and metabolic plasticity in AML ^30^. This selective vulnerability parallels prior work demonstrating that leukemic stem and progenitor cells rely on oxidative metabolism and exhibit limited capacity for glycolytic compensation ^5,6,50^. Unlike BCL-2 inhibition, which primarily exploits apoptotic priming, Clever-1 blockade appears to destabilize mitochondrial integrity by restricting lipid-supported membrane organization, suggesting a complementary mechanism that could be leveraged in combination strategies targeting mitochondrial metabolism.

Mitochondrial dysfunction induced by Clever-1 blockade was modest under nutrient-rich conditions but became pronounced under lipid-poor or physiologically relevant serum conditions. Adult human serum lipoproteins are enriched in unsaturated fatty acids compared with the more saturated lipid composition of fetal bovine serum, and serum lot variability can substantially influence culture outcomes ^51–53^. These differences likely affect intracellular lipid routing and mitochondrial membrane composition, providing a plausible explanation for why Clever-1–dependent mitochondrial fragility is masked under standard FCS conditions but revealed under human serum or lipid-restricted environments.

In summary, our data support a model in which Clever-1 contributes to metabolic robustness in AML by regulating intracellular lipid handling required for mitochondrial function. While bexmarilimab treatment alone does not reduce AML cell viability in vitro ^21^, Clever-1 blockade perturbs mitochondrial metabolism in a manner that lowers the adaptive capacity of leukemic cells. This provides a mechanistic framework for the previously observed sensitization of AML to standard-of-care therapies following Clever-1 targeting ^25^. Together, these findings identify Clever-1–dependent lipid handling as a metabolic vulnerability in AML and support the rationale for combining Clever-1–directed approaches with therapies that impose metabolic or mitochondrial stress.

## Acknowledgements

We thank Riikka Sjöroos, Mari Parsama and Teija Kanasuo for excellent technical support. We also thank the Laboratory of Electron Microscopy at the Institute of Biomedicine, University of Turku, for providing facilities and technical assistance in tissue sample preparation. Mass spectrometry analyses were performed at the Turku Proteomics Facility, supported by Biocenter Finland. This study was further supported by the Cell Imaging and Cytometry Core at the University of Turku and Åbo Akademi, and by Biocenter Finland. This study was funded by the Research Council of Finland and Sigrid Jusélius Foundation (MH).

## Authors’ Contributions

AY was responsible for conceptualization, methodology, investigation, data curation, software, formal analysis, visualization, and writing of the original draft, as well as writing–review and editing. JM and MiH contributed to investigation, data curation, formal analysis, and visualization. RT contributed to investigation and supervision. ST contributed to conceptualization, methodology, investigation, data curation, formal analysis, visualization, supervision, and writing–review and editing. MH contributed to conceptualization, methodology, formal analysis, visualization, supervision, funding acquisition, project administration, and writing–review and editing. All authors reviewed and approved the final version of the manuscript.

## Data Availability Statement

Data generated during the current study are available from the corresponding author on reasonable request.

