## Supplemental data for "Clever-1 blockade disrupts lipid metabolism and mitochondrial fitness in acute myeloid leukemia"

**Supplementary Materials, Ylitalo et al.**

**Legend for Supplementary Data File. Mass spec proteomics data from Clever-1 pulldowns after bexmarilimab treatment.**

The abundance of each identified protein from the 9-11 pulldowns of Clever-1 was divided by the abundance for the rat isotype control pulldown to give a "quotient" ratio. Any protein for which the quotient was less than 4 was not taken forward for further analysis. This filtering was sufficient to eliminate most common contaminants such as keratin.

The filtered hits for the hIgG4 treated KG-1 cells and the bexmarilimab treated KG-1 cells were then compared. The "difference in abundance" was calculated by subtracting the abundance for a protein pulled down by 9-11 from bexmarilimab treated cells (with rat isotype control pulldown abundance already subtracted) from the abundance of the same protein pulled down by 9-11 from hIgG4 treated cells (with rat isotype control pulldown abundance already subtracted). The "purity sum" for a protein was calculated by adding the quotient for the protein pulled down from bexmarilimab treated cells to the quotient for the protein pulled down from hIgG4 treated cells. The CRAPome database score for each protein was also retrieved.

The processed data is attached in an Excel file. The data in the tables are presented with red-to-blue heatmap coloring, with the greatest difference in abundance ranked at the top of the table. The two tables presented in the supplementary file list the proteins recovered less with Clever-1 after bexmarilimab treatment, and conversely the proteins recovered more with Clever-1 after bexmarilimab treatment

(from Figure 3B)

anti-Clever-1

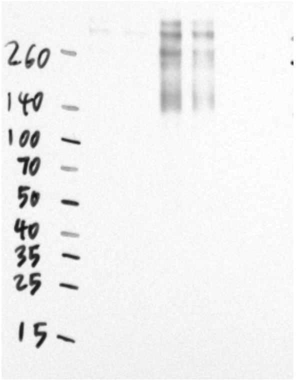

anti-ATAD3

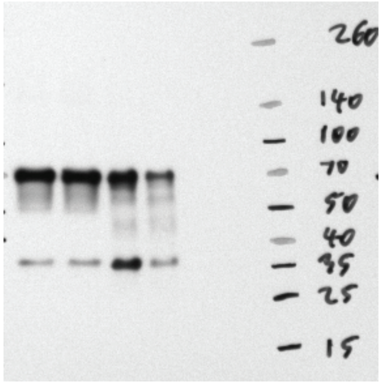

anti-IMPDH2

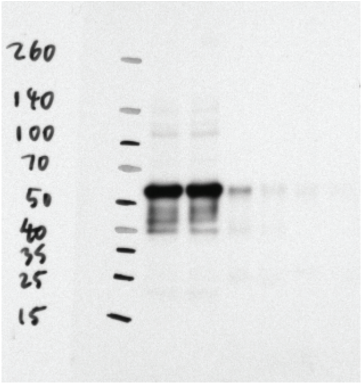

(from Figure 3C)

anti-Clever-1

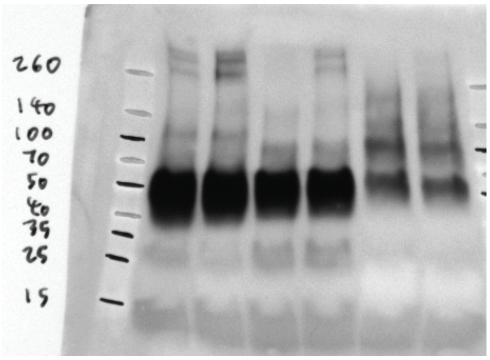

(from Figure 3D)

anti-Clever-1

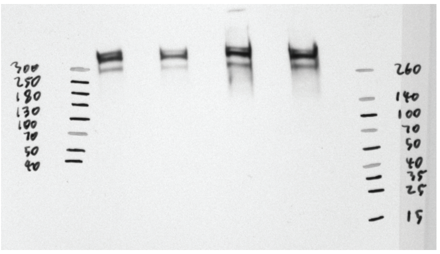

anti-ATAD3

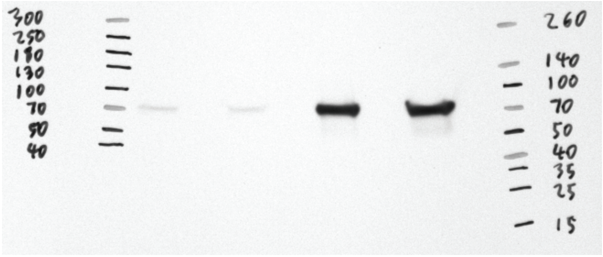

anti-PD1

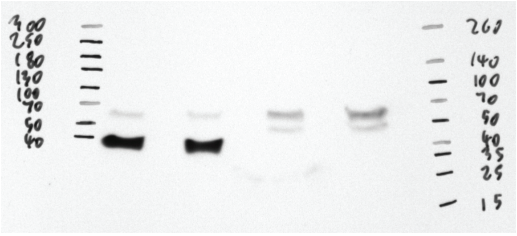

anti-GAPDH

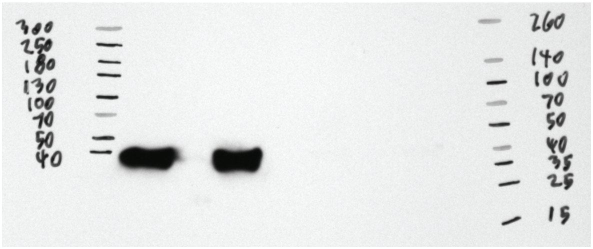

**Supplementary Figure 1.** Uncropped Western blots films from Figure 3.

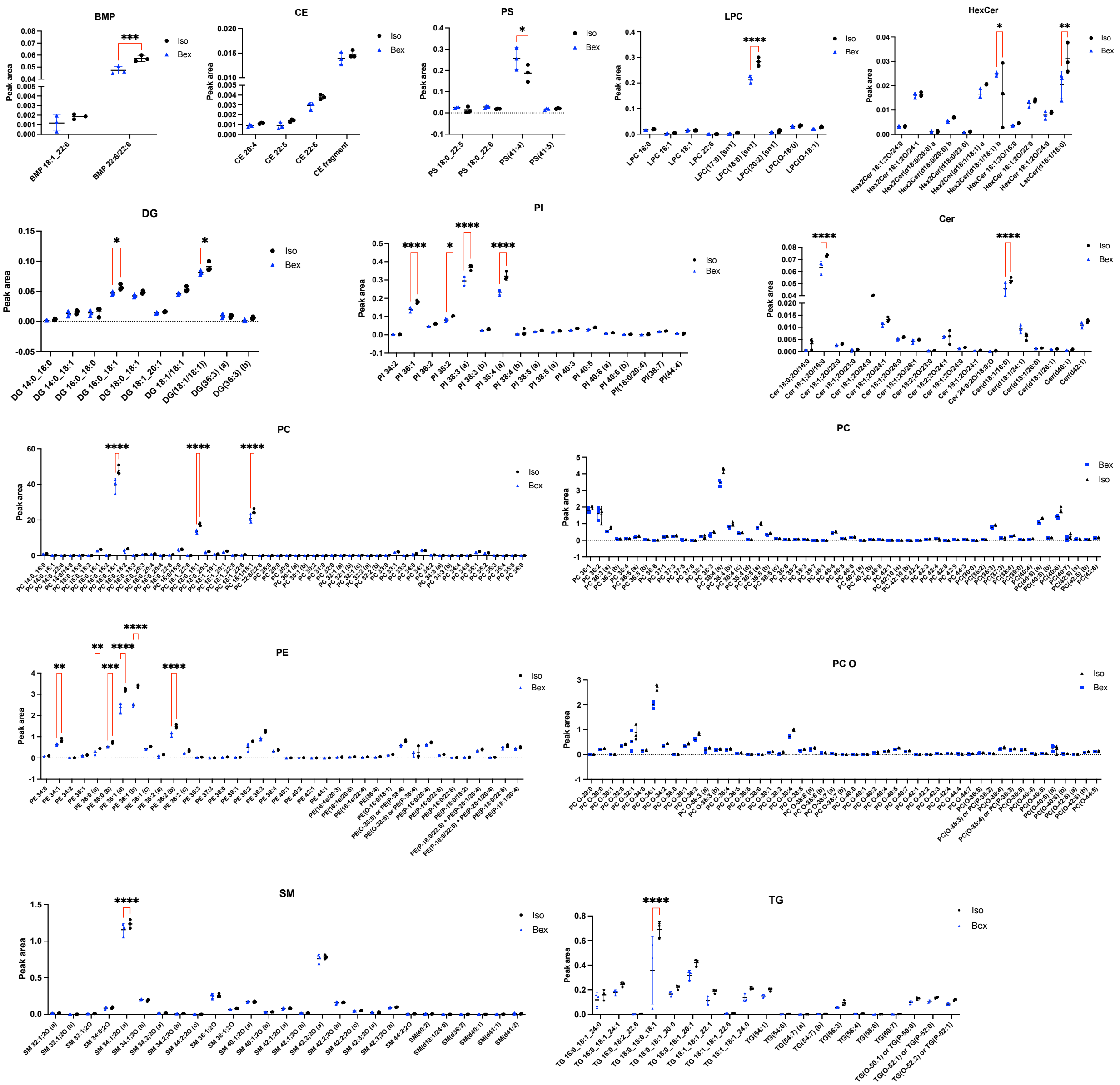


**Supplementary Figure 2. Data for individual lipid species in mitochondria following treatment.** KG-1 cells were cultured with hIgG4 isotype control or bexmarilimab, as described in the materials and methods. The mitochondria were isolated by MACS, and the mitochondrial lipids extracted for lipidomics analysis by mass spec. Experiments were repeated in triplicate. The plots summarize the peak area data for individual lipid species with grouped mean and SD. Statistical analyses was by 2-way ANOVA. BMP = bis(monoacylglycero)phosphates, CE = cholesteryl esters, Cer = ceramides, DG = diacylglycerols, Hex(2)Cer = dihexosylceramides, LPC = lysophosphatidylcholines, PI = phosphatidylinositols, PS = phosphatidylserines, SM = sphingomyelins, TG = triglycerides.

A

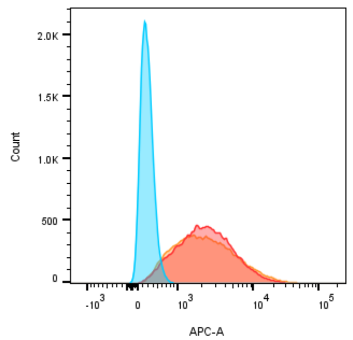


| Sample | Geometric MFI |
| --- | --- |
| Secondaries only | 285 |
| hIgG4 Isotype treated | 2208 |
| Bexmarilimab treated | 2215 |

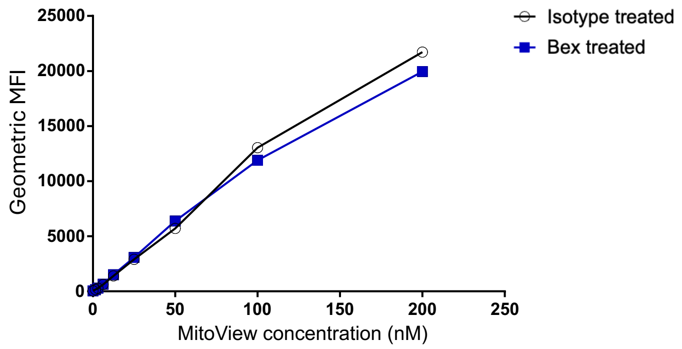


B

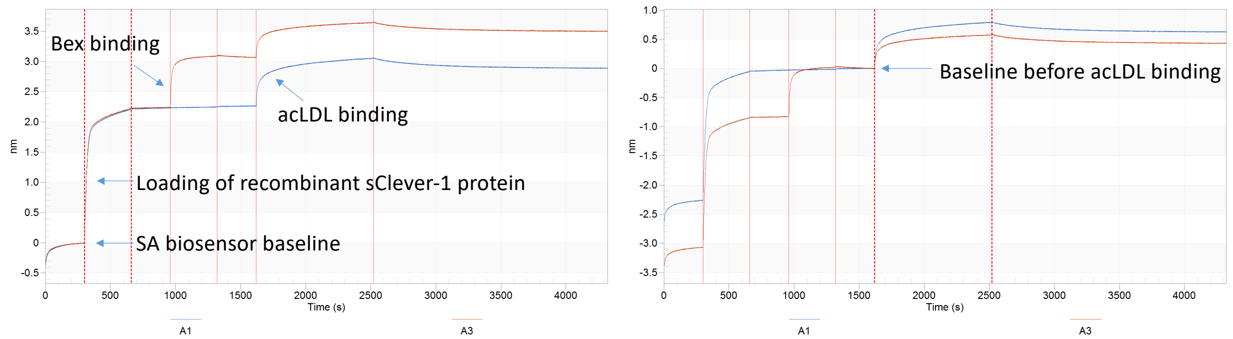


**Supplementary Figure 3. Bexmarilimab treatment does not directly prevent the interaction between Clever-1 and acLDL. A.** After bexmarilimab and acLDL treatment the relative quantity of live mitochondria within cells and the intensity of Clever-1 expression at the cell surface was measured. KG-1 were cultured with hIgG4 or bexmarilimab as described in materials and methods. For the final two hours they were cultured with A488-acLDL (Invitrogen) and for the final hour 1/16000 MitoView Fix(able) 640 far-red dye (Biotium) to label live mitochondria. Controls were prepared. The cells were stained with 9-11 antibody and analyzed via FACS. Geometric mean fluorescence intensity data is presented. **B.** The ability of bexmarilimab to directly block the binding of Clever-1 to acLDL was tested via Octet. Recombinant soluble secreted sClever-1 protein was produced and biotinylated as previously (Prince et al. 2025). The protein was loaded onto streptavidin SA biosensors on Octet 384 Red (Forte Bio). One sClever-1 loaded biosensor was saturated with bexmarilimab, and the other with hIgG4 isotype control. Both sensors were then dipped into acLDL (Thermo Fisher Scientific) in PBS + 0.2% BSA for association followed by dissociation. The BLI data was analyzed and aligned using Octet Data Analysis software. The aligned sensorgrams are presented.
